# A Novel Virus Discovered in the Yeast *Pichia membranifaciens*

**DOI:** 10.1101/2022.01.05.475065

**Authors:** Mark D. Lee, Jack W. Creagh, Lance R. Fredericks, Angela M. Crabtree, Jagdish Suresh Patel, Paul A. Rowley

## Abstract

Mycoviruses are widely distributed across fungi, including yeasts of the Saccharomycotina subphylum. It was recently discovered that the yeast species *Pichia membranifaciens* contained double stranded RNAs (dsRNAs) that were predicted to be of viral origin. The fully sequenced dsRNA is 4,578 bp in length, with RNA secondary structures similar to the packaging, replication, and frameshift signals of totiviruses of the family *Totiviridae.* This novel virus has been named Pichia membranifaciens virus L-A (PmV-L-A) and is related to other totiviruses previously described within the Saccharomycotina yeasts. PmV-L-A is part of a monophyletic subgroup within the I-A totiviruses, implying a common ancestry between mycoviruses isolated from the *Pichiaceae* and *Saccharomycetaceae* yeasts. Energy minimized AlphaFold2 molecular models of the PmV-L-A Gag protein revealed structural conservation with the previously solved structure of the *Saccharomyces cerevisiae* virus L-A (ScV-L-A) Gag protein. The predicted tertiary structure of the PmV-L-A Pol and its homologs provide details of the potential mechanism of totivirus RNA-dependent RNA polymerases (RdRps) because of structural similarities to the RdRps of mammalian dsRNA viruses. Insights into the structure, function, and evolution of totiviruses gained from yeasts is important because of their parallels with mammalian viruses and the emerging role of totiviruses in animal disease.

## Introduction

*Pichia membranifaciens* is a pseudofilamentous fungi and is a common spoilage organism, contaminating meats, soft cheeses, vegetables, and beverages [1]. Although *P. membranifaciens* is frequently associated with spoilage, it can act as a biocontrol agent for both plant pathogens and aggressive post-harvest spoilage organisms [2–8]. Biological control by *Pichia* species relies on the production of volatile organic compounds, killer toxins, and glucanases [9–12]. Killer toxin production by yeast is often associated with the presence of extrachromosomal genetic elements, including double-stranded RNAs (dsRNAs). Previous attempts to discover novel dsRNA-associated killer toxins found that *P. membranifaciens* strain NCYC333 plays host to a putative dsRNA mycovirus, most likely of the family *Totiviridae* [9,13]. The number of mycoviruses has expanded over recent years, with the International Committee for the Taxonomy of Viruses (ICTV) recognizing mycoviruses with both RNA and DNA genomes, with the majority being linear dsRNAs [14]. Mycoviruses can be found across all major taxonomic groups of fungi and are not thought to spread via extracellular transmission, instead transmitting vertically by mitosis and meiosis, and horizontally due to anastomosis or yeast mating. Generally, mycoviral infections are considered benign with little obvious fitness cost to the host. However, there are instances where mycovirus infection results in noticeable phenotypes, such as the alteration of fungal virulence, morphology, growth rate, and pigmentation, and so have been used to control fungal diseases [15,16]. To date, there have been no descriptions of mycoviruses in *P. membranifaciens.* Given that mycovirus infection can alter host physiology and *P. membranifaciens* holds future promise as a biological control agent, it is important to better characterize the viruses associated with the species.

Mycoviruses in the family *Totiviridae* are dsRNA viruses that are recognized as being distributed across diverse fungal species *(Totivirus* and *Victorivirus* genera) but are also found in species of human and animal pathogenic protozoa *(Giardiavirus, Trichomonasvirus,* and *Leishmaniavirus* genera) [17–21]. In addition, there are reports of totivirus-like viruses that infect shrimps, and mosquitos [22,23]. A well-studied member of the *Totiviridae* family is the L-A virus of the brewer’s/baker’s yeast *Saccharomyces cerevisiae* (ScV-L-A; Saccharomyces cerevisiae virus LA) [20,24]. This virus has a dsRNA genome of 4,580 bp and encodes two proteins, Gag and Gag-pol, and replicates within the cytosol of the yeast cell. Gag is responsible for the encapsidation of the dsRNA genome of the totivirus and for the binding and decapping of host mRNAs, which provides newly synthesized viral transcripts with 5’ cap structures [25]. A frameshift is required for the translation of Gag-pol and requires a stem-loop and slippery site to induce ribosome stalling and −1 frameshifting [26]. The Gag-pol fusion enables the tethering of the Pol domain to the inner surface of the viral capsid (1-2 per capsid). The Pol domain is an RNA-dependent RNA polymerase (RdRp) of unknown structure that is essential for (+) sense RNA packaging and the synthesis of complementary (-) sense RNAs to form the dsRNA genome [27]. The genomic dsRNA is used as a template for conservative RNA replication to produce (+) sense viral transcripts that are extruded to the cytosol for translation and packaging into new viral particles.

In this manuscript we describe the discovery and characterization of a novel member of the *Totiviridae* isolated from *P. membranifaciens.* As the first example of a dsRNA virus in the species we have named it Pichia membranifaciens virus L-A (PmV-L-A), consistent with the naming of the first yeast virus discovered in *S. cerevisiae.* The organization of the genome is the same as other related yeast totiviruses, with conserved RNA structures that are likely important for RNA frameshifting, replication, and packaging. The nucleotide and protein divergence would classify this virus as a new species and is an outgroup to other viruses of the *Saccharomycetaceae*. Structural modeling of the PmV-L-A proteins are consistent with known totivirus capsid proteins and RdRp proteins of dsRNA viruses that infect mammals.

## Methods

### Purification and digestion of dsRNAs with RNAse III

Double-stranded RNAs were purified from *P. membranifaciens* strain NCYC777 (obtained from the National Collection of Yeast Cultures, UK) using the method described previously. [13]. Five μg of dsRNAs were incubated with 10 units of ShortCut® RNase III (New England Biolabs) in the buffer conditions recommended by the manufacturer for 20 min at 37°C. The reaction was stopped by the addition of EDTA to a final concentration of 45 mM. DNase I (New England Biolabs) digestion was performed as directed by the manufacturer in 1X reaction buffer for 10 min at 37°C. the reaction was stopped by incubation at 75°C for 10 min. All digested products were analyzed by agarose gel electrophoresis with ethidium bromide staining.

### The determination of the genetic sequence of dsRNAs

All methods for the sequencing of dsRNAs were as previously described [13]. 5’ RACE was carried out as described by the manufacturer’s instructions using purified dsRNAs (Thermo Fisher catalogue #18374058). Amplification of the 5’ ends of the dsRNA using RACE used the specific primers GSP1: 5’-ACACCATTGTTAGTACG-3’ and 5’-GAATATACCAGTTGAGG- 3’ and GSP2: 5’-GAAGATGATCCACCAACAATAACAGG-3’ and 5’- AGTGGGAAAGGGCAATGTATGG-3’. Amplification of cDNAs from PmV-L-A dsRNAs by RT-PCR was carried out with the following primer combinations in Figure 2: Reaction 1: 5’- TCCAGTCAATGCTGATAGAGG-3’ and 5’-AGCGGAGCTTCAATACCTGA-3’. Reaction 2: 5’-GCTTATTCAGAGGGGTGGTG-3’ and 5’-AACTGAGCCACCCGAGAATA-3’. Reaction 3: 5’-CGTGGCAGTCAAAAGAAA-3’ and 5’-AACTGAGTCGCACACCCAAT-3’. Reaction 4: 5’-TCGCGATGTTTAGGTGTGAA-3’ and 5’-GTGGTATGCCGACGAATTTT-3’.

### Phylogenetic analysis of PmV-L-A

Gag and Pol protein sequences from PmV-L-A and its homologs were aligned using MUSCLE and inspected manually for accuracy. MEGAX was used to determine the appropriate amino acid substitution matrix for each alignment dataset. MEGAX and PhyML were used to create phylogenetic models of the evolutionary relationship between the different amino acid sequences [28,29]. For the MEGAX analysis the maximum likelihood and neighbor joining methods were used with 100 bootstrap replicates. PhyML parameters included having the program estimate the proportion of invariable sites from the aligned dataset, using BEST tree topology estimation, a random starting tree, and 100 bootstrap replicates. The totivirus Saccharomyces cerevisiae virus L-BC (ScV-L-BC) was used as an outgroup in all phylogenetic models.

### Molecular modeling of the Gag and Pol proteins of PmV-L-A

AlphaFold2 version 1.2 was used to build 3-D protein structure models using the amino acid sequences derived from the *gag* and *pol* genes [30]. Amino acid sequence was used as an input with default AlphaFold2 model building parameters (msa_mode: MMseqs2 (uniref+environment); model_type: auto; pair_mode: unpaired+paired; num_recyles: 3). Five Gag models were generated with similar local distance difference test (LDDT) values per residue (Figure S2) while AlphaFold2 could only predict one model for Pol proteins because of their length and low similarity. This resulted in the best predicted protein model for each PmV-L-A Gag, PmV-L-A Pol, ScV-L-A Pol, TdV-LABarr1 Pol, and TAV-1 Pol amino acid sequences, which were then subjected to energy minimization. Prior to energy minimizations, the catalytic site Mg^2+^ ion of each RdRp structure was identified and transferred to predicted model structure by rigid-body backbone alignment of the relevant region of the crystal structure of RNA dependent RNA polymerase domain of west Nile virus (PDBID: 2HFZ) using Pymol visualization software package. Energy minimization was carried out for each model using standard energy minimization protocol described in our previous study [31]. Briefly, each model was placed in a dodecahedron solvent box, and a 10 Å TIP3P water layer was added to solvate the protein model. Na+ and Cl- ions at a concentration of 0.15 mol/L were then added to the water layers to maintain the charge neutrality. The AMBER99SB*-ILDNP force field parameters were used for the protein and ions. Each system was subjected to energy minimization using the steepest descent algorithm for 10,000 steps using the GROMACS package [32]. Stereochemical checks were performed for each model after the energy minimization using the SWISS-MODEL structure assessment tool (https://swissmodel.expasy.org/) to assess the quality of the predicted structures. The final energy minimized Gag and Pol model structures were analyzed and compared to related viral proteins using the Pymol visualization software package.

## Results

### Purification and Digestion of Double-Stranded RNAs from P. membranifaciens

The previous identification of *P. membranifaciens* killer yeasts had resulted in the screening of different strains for the presence of totiviruses and M satellite dsRNAs using cellulose chromatography [13]. This approach identified a putative dsRNA species within *P. membranifaciens* strain NCYC333 (syn. ATCC 36908, CBS 7374) that was isolated from draught beer in the United Kingdom and deposited in the National Collection of Yeast Cultures in 1953 [12]. The dsRNA had an electrophoretic mobility that was suggestive of a totivirus (Figure 1, lane 1). To confirm that this discrete band was either dsRNA or DNA, the purified nucleic acids were incubated with DNase I or RNase III. The nucleic acid was completely digested in the presence of RNase III (Figure 1, lane 2).

**Figure 1.**
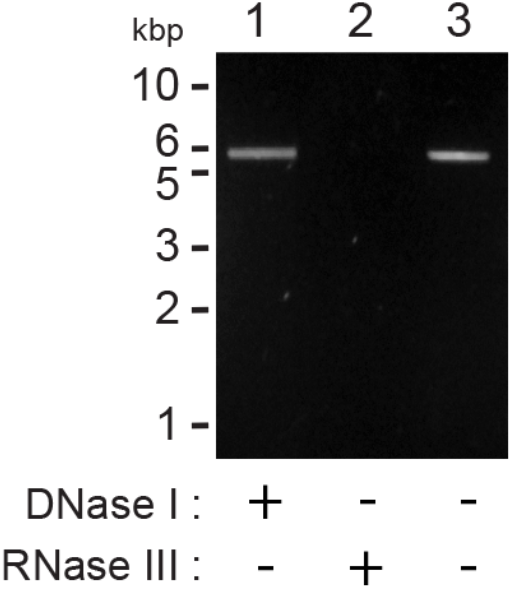
Extrachromosomal nucleic acids within *P. membranifaciens* are double stranded RNA molecules. Agarose gel electrophoresis of cellulose column purified nucleic acids from *P. membranifaciens* NCYC333 with DNase I (lane 1) and ShortCut® RNase III (lane 2).

### Determining the Nucleic Acid Sequence of the Double-Stranded RNA from P. membranifaciens

With the confirmation that *P. membranifaciens* NCYC333 was host to dsRNAs of unknown identity, purified dsRNAs were subjected to poly(A) tailing, cDNA creation, and Illumina shortread DNA sequencing as described previously [13]. Assembly and analysis revealed that the dsRNA species was the genome of a totivirus similar to those that have been previously described in yeasts of the *Saccharomycetaceae* (Figure 2A). Specifically, the assembly of 42,960 polished Illumina sequence reads revealed five contigs that were more than 1,000 bp in length. Blastx analysis identified that four of these contigs were related to yeast totiviruses (Figure 2B, red dots). After assembly into a contiguous nucleotide sequence, primer pairs were designed to confirm this organization by reverse-transcriptase PCR (RT-PCR) and Sanger sequencing (Figure 2C). The terminal ends of the dsRNA were determined using 5’ RACE and confirmed the total length of the dsRNA was 4578 bp, which is consistent with a totivirus genome (Genbank accession number OL687555). Analysis of the complete dsRNA sequence identified an open reading frame of 680 amino acids from nucleotide 28 to 2070 (Figure 2A). Blastp of this amino acid sequence revealed that it is most closely related to the Gag protein of totiviruses found with other species of budding yeasts from the *Pichiaceae* and *Saccharomycetaceae* such as Ambrosiozyma totivirus A (AkV-A; 56% identity), Saccharomyces cerevisiae virus L-A (ScV-L-A; 41% identity) and Torulaspora delbrueckii virus L-A (TdV-L-A; 45% identity). The Gag protein also contained a conserved histidine at position 154 that has been shown to be critical for cap snatching activity of ScV-L-A [25,33]. Analysis of the 3’ end of the *gag* gene revealed the sequence of a −1 ribosomal frameshift site with a stem loop downstream of a 5’- GGGUUU-3’ slippery sequence (Figure 2D). This would enable the creation of a Gag-Pol fusion protein of 1522 amino acids that is 49% and 50% identical to ScV-L-A and TdV-L-A, respectively (Figure 2D). Other characteristic RNA secondary structures were also evident at the 3’ terminal end of the RNA with stem loops characteristic of totivirus replication and packaging signals (Figure 2A). The latter contained the typical 5’ A-bulge and an 11-nucleotide stem loop observed in other totiviruses and satellite dsRNAs (Figure 2D) [34]. As the first dsRNA virus of the species, we named the new virus Pichia membranifaciens virus L-A (PmV-L-A).

**Figure 2.**
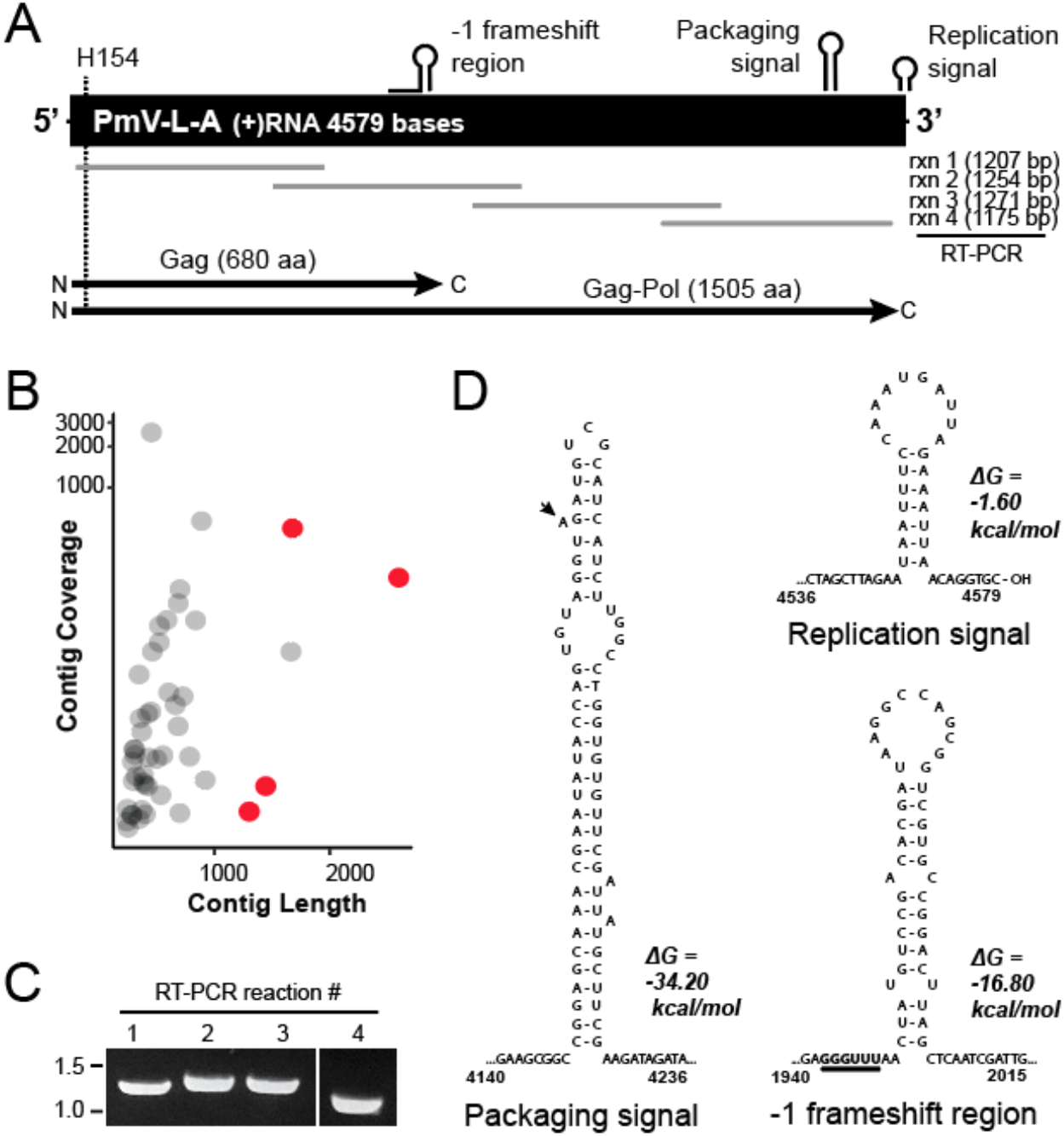
A dsRNA virus within *P. membranifaciens* resembles the genomic organization of a totivirus. (A) Schematic overview of the organization of PmV-L-A genome including the position of RNA secondary structure, ORFs, and the conserved H154 required for cap snatching activity of Gag. Rxn1-4 show the position of RT-PCR products that were used to confirm the genome sequence of PmV-L-A by Sanger sequencing. (B) Sequence contigs after *de novo* assembled represented by contig coverage and contig length. Blastx analysis of the ten contigs with the longest length enabled the identification of four sequences related to yeast totiviruses (red). (C) Products of RT-PCR using overlapping primer pairs to confirm the assembled sequence of PmV-L-A. (D) RNA secondary structure predictions by mFold [35].

### Phylogenetic Analysis of PmV-L-A

To determine the evolutionary relationship of PmV-L-A to other previously described totiviruses, blastp was used to identify homologs of PmV-L-A Gag and Pol proteins. The search results found that the homologous proteins were from viruses of the genus *Totivirus* in subgroup I-A. Results were filtered to only include those proteins that were >90% the length of the PmV-L-A proteins, and multiple sequence alignments were created using MUSCLE. The Gag and Pol proteins from the subgroup I-B totivirus ScV-L-BC were used as outgroup proteins in the alignments. The alignments were used as inputs for phylogenic analysis using neighbor joining and maximum likelihood methodologies implemented by MEGAX and PhyML [28,29]. In all phylogenic models PmV-L-A Gag and Pol were outgroups to the totiviruses found within the *Saccharomycetaceae* (Figure 3 and S1), consistent with *P. membranifaciens* being from the family *Pichiaceae* [36]. PmV-L-A Gag was most closely related to the Gag protein from the incomplete genome of Ambrosiozyma totivirus A identified within the yeast *Ambrosiozyma kashinagicola* isolated in Japan in 2016 (Genbank accession number MK231133.1). Both *P. membranifaciens* and *A. kashinagicola* have been classified within the *Pichiaceae* family. PmV-L-A proteins appeared to share a common ancestry with another fungal totivirus (tuber aestivum virus 1) and a hypothetical protein in the genome of the ascomycetous yeast *Scheffersomyces stipitis* of the CUG yeast clade that includes important human pathogens such as *Candida auris* and *Candida albicans* [36]. Together, PmV-L-A proteins and their homologs formed a monophyletic group that are separate from the other subgroup I-A totiviruses that are associated with both fungi and plants (Figure 3).

**Figure 3.**
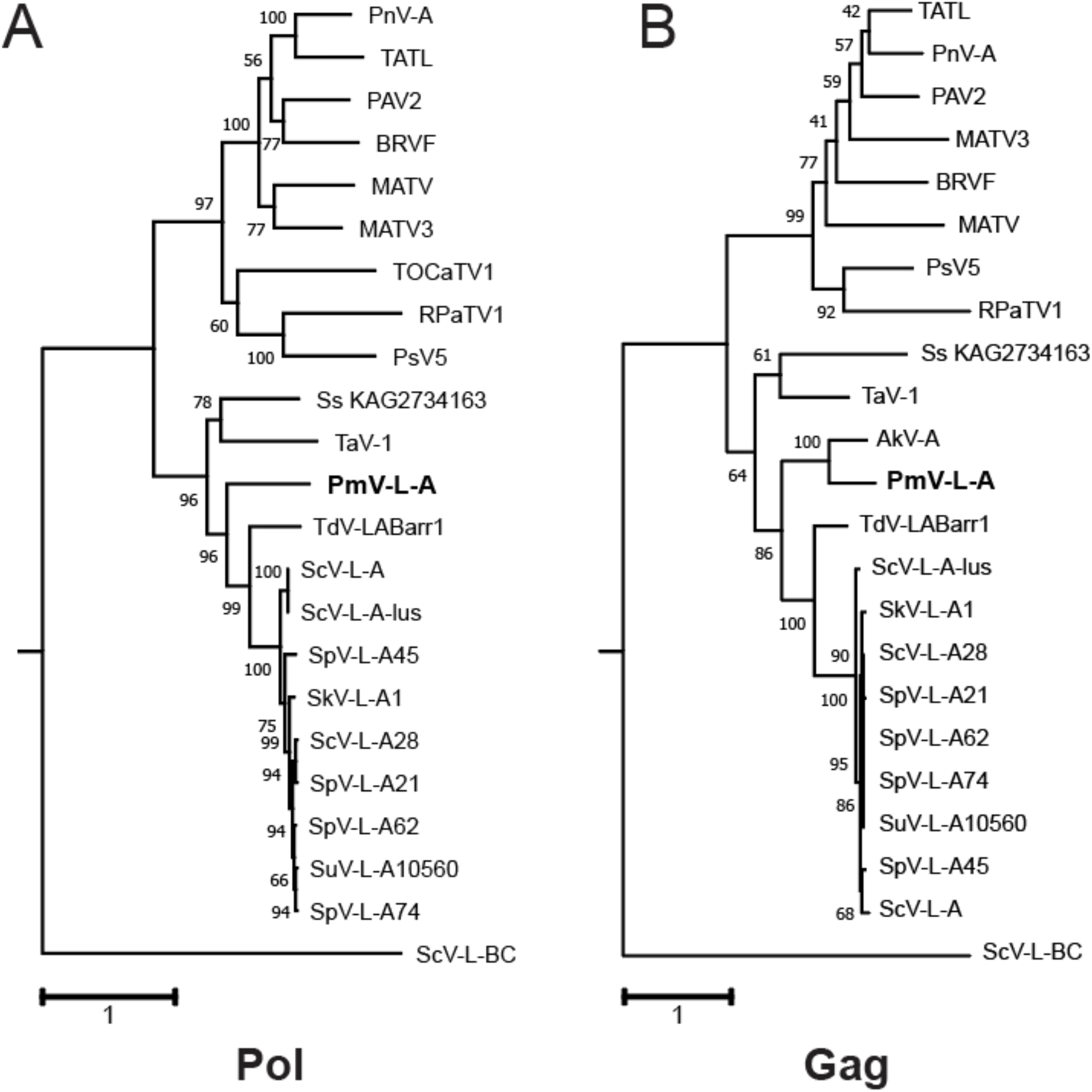
Phylogenetic analysis of the Gag and Pol proteins encoded by PmV-L-A. Rooted maximum likelihood phylogenic models were constructed using PhyML from the amino acid sequences of the (A) Pol and (B) Gag proteins of PmV-L-A and related proteins identified by blastp. The LG model with a gamma distribution (G) and invariable sites (I) was determined to be the substitution matrix that fit best with these datasets. The numbers at each node are the bootstrap values from 100 iterations. The scale bar represents the distance of one amino acid substitution per site. The amino acid sequences in the phylogeny are derived from viruses listed in Table S1.

### Structural Modeling of the Gag and Pol Proteins from PmV-L-A

AlphaFold2 was used to predict 3-D structural models to determine whether the Gag and Pol proteins from PmV-L-A were similar in tertiary structure to models of related virus proteins [30]. AlphaFold2 predicted five 3-D models for Gag amino acid sequence while it could only predict a single 3-D model for each Pol amino acid sequence. The LDDT per residue score was used to select the best model out of the five predicted Gag models. However, the LDDT score was found to be highly similar for each Gag model and therefore the first model was selected for further assessment (Figure S2). After generating the models, structures were subjected to energy minimization using GROMACS software package. Energy minimized structural models were then used as inputs to carry out stereochemical checks to assess the quality of the predicted models (Figure S2). Molprobity scores, which combine the clashscore, rotamer, and Ramachandran evaluations into a single score was less than 1.4 Å for all computational models, which suggests good overall quality of the predicted and energy minimized 3-D structural models (Table S2).

Final selected Gag model of the PmV-L-A Gag protein was highly similar in structure to the crystal structure model of ScV-L-A Gag (PDB: 1M1C) [37]. Overlaying the two structures revealed a root mean squared deviation (RMSD) of only 1.7 Å using PDBeFold. In addition, 91% of the secondary structure of the ScV-L-A Gag was identified in the model of PmV-L-A, despite the amino acid identity of only 41%. The modeled Gag also had 95% of amino acid residues in the Ramachandran favored region (Figure S2). In both models, the catalytic histidine 154 was also conserved and positioned on the outer surface of Gag located at the tip of a surface trench formed from four loops (Figure 4A).

**Figure 4.**
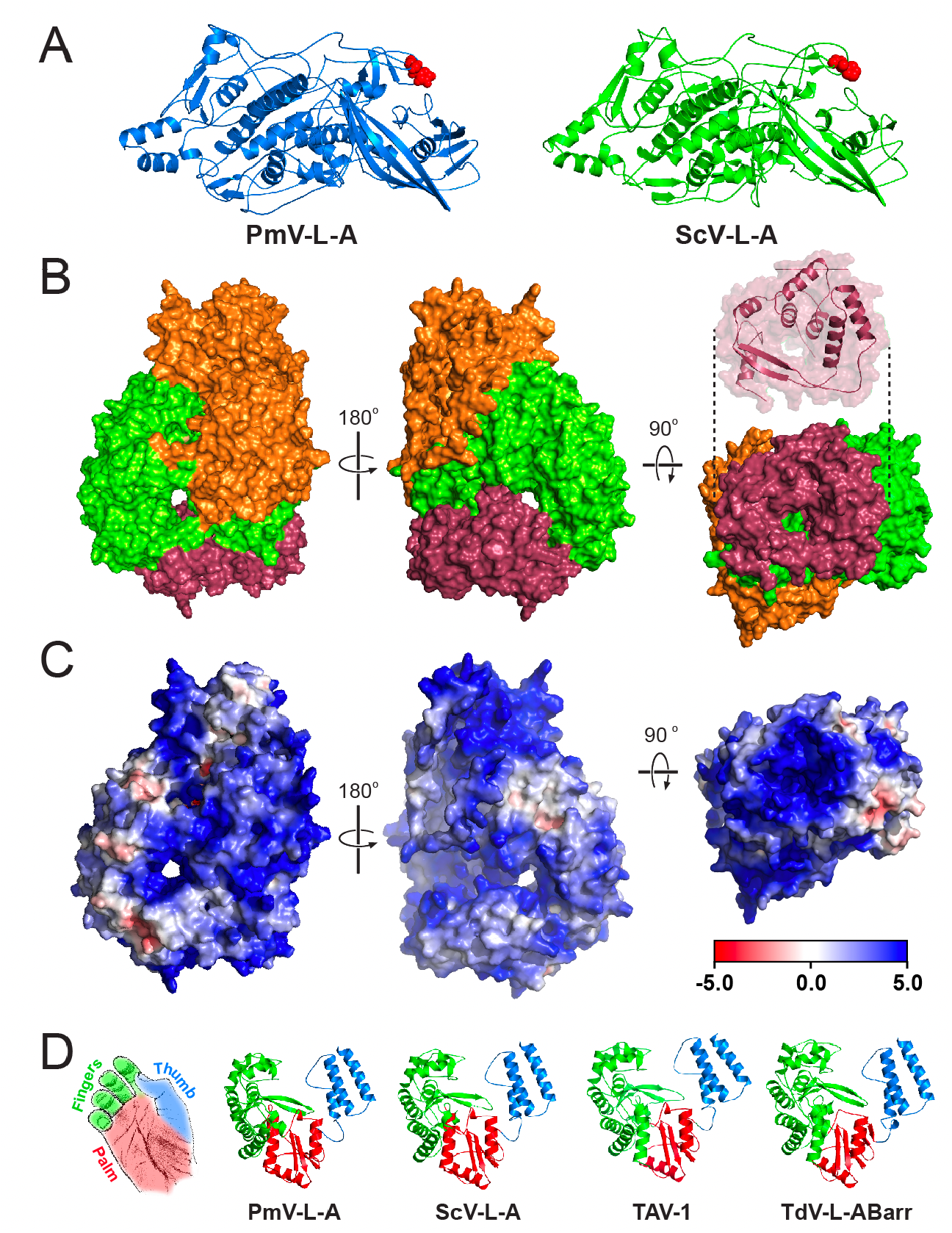
Structural models of the PmV-L-A Gag and Pol proteins and comparison to other totiviral polymerases. AlphaFold2 modeling server was used to generate molecular models of proteins encoded by PmV-L-A with energy minimization using GROMACS software. (A) Cartoon models of the Gag protein of PmV-L-A compared to the crystal structure model of the Gag protein of ScV-L-A. The catalytic histidine (H154) is depicted as red spheres in each model. (B) Surface model of PmV-L-A Pol protein depicting the N-terminal domain (orange), catalytic domain (green), and bracelet domain (burgundy). The bracelet domain is also depicted as a cartoon to better illustrate the channel structure. (C) Surface electrostatic model of the PmV-L-A Pol using adaptive Poisson Boltzmann solver (APBS). (D) Cartoon representation of the catalytic domain of four totivirus polymerases illustrating the finger, thumb, and palm subdomains.

Unlike Gag, there are no structural models of totivirus polymerases and there is <20% sequence identity to other RNA-dependent RNA polymerases (RdRp) that have been crystalized. The modeled structure of the Pol proteins of PmV-L-A, ScV-L-A, TdV-LABarr1, and TAV-1 were created using the same methods as for Gag, and the predicted RdRp structures were found to match the structures of other viral RdRp proteins using PDBeFold. The longest structural match with PmV-L-A Pol was with VP1 of rotavirus SA11 (PDB 2R7T; group III dsRNA virus, family *Reoviridae* [38]) with 488/866 aligned amino acids, 9.2% identical residues, and an RMSD of 3.96 Å. Similar results were obtained for all other modeled totivirus polymerases. The modeled Pol proteins also had 90.20 – 92.89% of amino acid residues in the Ramachandran favored region (Figure S2). Using the crystal structure model of rotavirus VP1 as a guide, it was possible to identify three distinct domains in the totivirus polymerases. The core catalytic domain (amino acids 310-697) was clearly discernible sandwiched between an N-terminal domain (1-309), and a C-terminal ‘bracelet’ domain (698-869) (Figure 4B). The totivirus Pol proteins all contained a central cavity with four entry channels that are positioned in a similar manner to those of VP1 (Figure 5C) [39]. The structure of the C-terminal bracelet domain of the totiviruses encircles one of these pores that in VP1 is used as an exit for the RNA templates during replication. In totiviruses, this domain is smaller (172 versus 392 (VP1) amino acids) with a single predicted antiparallel beta sheet and seven alpha helices compared to the complex VP1 bracelet domain of 20 alpha helices and two beta sheets. Electrostatic charge maps indicate that these channels are positively charged, with obvious tracks of basic residues that lead to these entry points (Figure 4C). The catalytic domain of the modeled RdRps adopted tertiary structures characteristic of a closed right-hand with palm, fingers, and thumb subdomains (Figure 4D). As with VP1, the palm was an antiparallel β sheet supported by three α helices and the thumb consisting of three α helices that interacted with the loops of the fingers and enclosed the central cavity. These subdomains contained seven motifs (from the N-terminus, G, F, A, B, C, D, and E) with conserved residues that are important for RNA polymerization by RdRp proteins. Our models confirm that these motifs are exposed within the central cavity of the RdRp. Motif F, and G formed part of the fingers subdomain, with motif F interacting with motif E of the palm subdomain to close the catalytic domain. The remaining motifs were found in the palm subdomain (motif A, B, C, and D) and are positioned to facilitate catalysis, specifically through metal ion coordination (motifs A and C) and nucleotide triphosphate binding (motif B).

## Discussion

In this article we report the first identified species of dsRNA virus found in the yeast *P, membranifaciens*. The virus is of the *Totiviridae* family and although it has diverged from other previously described yeast totiviruses, there is sequence and structural conservation in both the viral RNAs and proteins. The relatedness of PmV-L-A to other yeast totiviruses is consistent with the relatedness of the different yeast host species. Specifically, the positioning of PmV-L-A as being most closely related to Ambrosiozyma totivirus A (AkV-A) that was identified within *Ambrosiozyma kashinagicola,* a member of the *Pichiaceae* and as an outgroup to totiviruses found in yeasts of the *Saccharomycetaceae* (i.e. ScV-L-A; *S. cerevisiae* and TdV-LABarr; *T. delbrueckii).* The relatedness of these different species of totiviruses implies that long-term coevolution of PmV-L-A with *P. membranifaciens.* This is consistent with yeast totiviruses being transmitted between hosts via mating and not via an extracellular mode of transmission. However, we have previously shown that other strains of *P. membranifaciens* do not host dsRNAs [13]. This suggests that these strains have either lost their totiviruses or that *P. membranifaciens* NCYC333 is unique in its acquisition of a totivirus. Interestingly, *P. membranifaciens* contains the genes that would constitute an active RNAi system, which might be expected to prevent virus infection [40,41]. It remains to be determined if the Dicer-like and Argonaute-like proteins of *P. membranifaciens* NCYC333 have been inactivated, either by mutation or by an undescribed totivirus-encoded countermeasure. Indeed, there are examples of mycoviruses that produce effectors that interfere with RNAi [42–44]. *P. membranifaciens* has potentially hosted totiviruses that are more closely related to viruses found within *Saccharomyces* yeasts as evinced by genome integrated viral sequences in *P. membranifaciens* [45]. This would suggest the possibility of horizontal transmission of viral dsRNAs between yeasts as has also been shown by phylogenic analysis of mycoviruses and satellite dsRNAs from different yeast species [13,45,46].

The positioning of RNA secondary structure elements in the genome of PmV-L-A suggests that they are functionally equivalent to those first described and characterized in ScV-L-A. Although there is only 54% nucleotide sequence identity between PmV-L-A and ScV-L-A, the similarity of RNA secondary structures indicates conserved strategies for packaging, replication, and frameshifting across yeast totiviruses. For example, the slippery sequence for the −1 frameshift in PmV-L-A is identical to that of ScV-L-A. Furthermore, the putative packaging signal of PmV-L-A is predicted to form a stem-loop with the essential A bulge and also the G and C nucleotides at the beginning and end of the loop that are important for encapsidation by ScV-L-A Pol [34]. The encapsidation of the ScV-L-A RNA transcripts is dependent on the first 213 amino acids of the Pol protein [47]. Our structural model shows that this N-terminal domain is distinct from the catalytic and bracelet domains and is likely important for stabilizing and enclosing the catalytic core of the polymerase. Consistent with its role in packaging, the N-terminal domain also has large surface tracks of positive charge that could help to bind viral RNAs. The computational modeling of the totivirus Pol proteins has also shown that the domain organization is consistent with the RdRp subdomains of the palm, fingers, and thumb of other RNA viruses, particularly the group III dsRNAs viruses of the *Reoviridiae* (including rotaviruses and reoviruses). The importance of conserved catalytic motifs A-G that are common to many RdRps has been previously confirmed for ScV-L-A Pol that we now show are positioned in a similar manner to other RdRps [48–50]. There are also additional conserved motifs specifically within totivirus RdRps that have been identified and named motif 1 and 2 [48]. In our models, motif 1 is important for contact with motif F in the fingers and motif 2 is within the fingers domain and exposed to the inner catalytic cavity close to motif D. As with the RdRps of the *Reoviridae,* the PmV-L-A Pol structural model is permeated by four positively charged pores that would enable strands of RNA to enter and exit the polymerase and the diffusion of small molecules and ions into the catalytic core. The structural similarity to the four-tunnel RdRps of *Reoviridae* would suggest that totivirus Pol proteins allow ingress and egress of RNAs (genomic dsRNAs and ssRNAs) in a similar manner.

## Acknowledgements

The authors would like to thank Lauren Saucedo for drawing of the image in Figure 4D. Research reported in this publication was supported by the National Institute of General Medical Sciences of the National Institutes of Health under Award Number P20GM104420 and the National Science Foundation Division of Molecular and Cellular Biosciences grant number 1818368. The content is solely the responsibility of the authors and does not necessarily represent the official views of the National Institutes of Health or the National Science Foundation.

---

**Figure S1.**
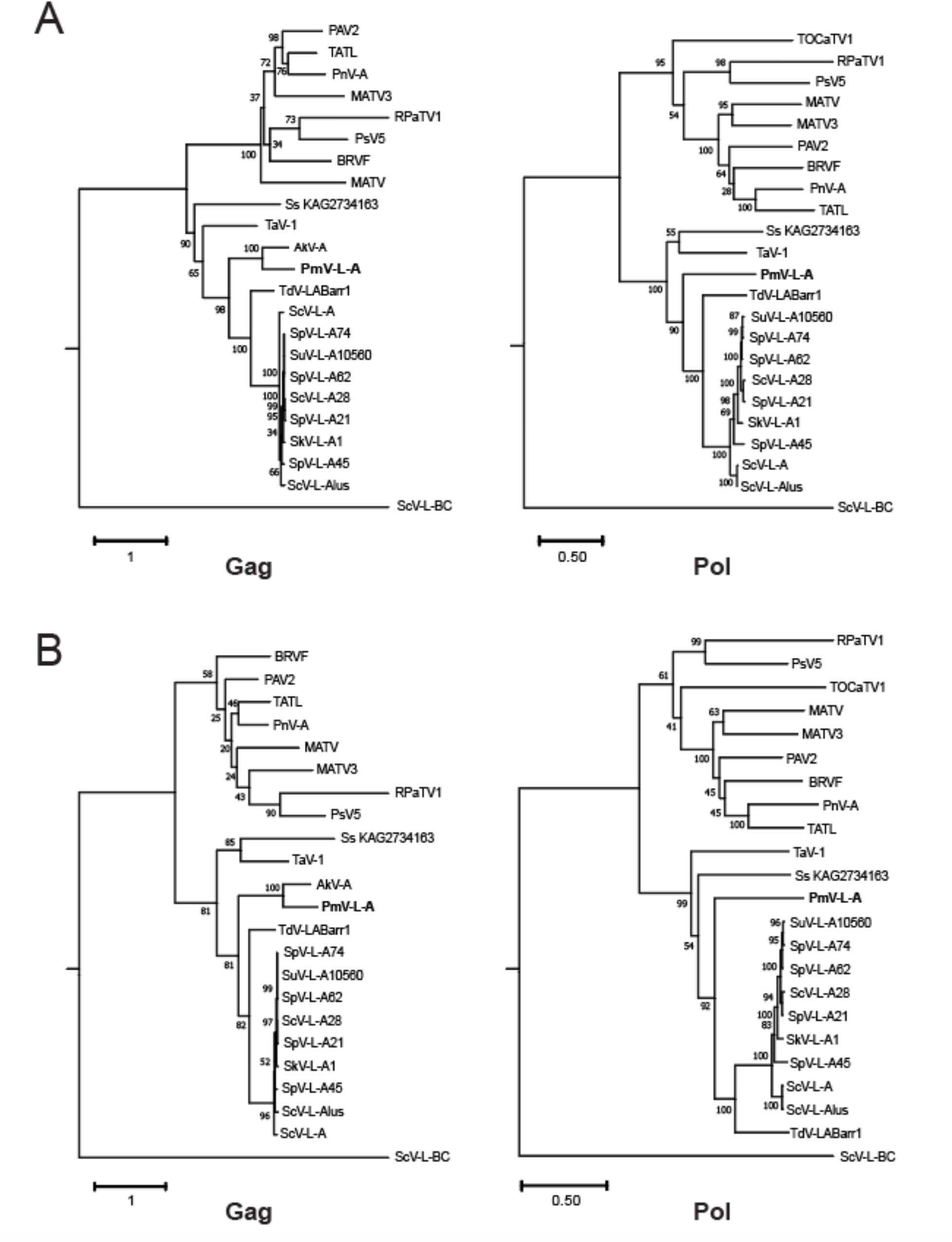
Phylogenetic analysis of the Gag and Pol proteins encoded by PmV-L-A. (A) Rooted maximum likelihood and (B) neighbor joining phylogenic models were constructed using MEGAX from the amino acid sequences of the Pol and Gag proteins of PmV-L-A and related proteins identified by Blastp. The LG model with a gamma distribution (G) and invariable sites (I) was determined to be the substitution matrix that fit best with these datasets. The numbers at each node are the bootstrap values from 100 iterations. The scale bar represents the distance of one amino acid substitution per site. The amino acid sequences in the phylogeny are derived from viruses listed in Table S1.

**Figure S2.**
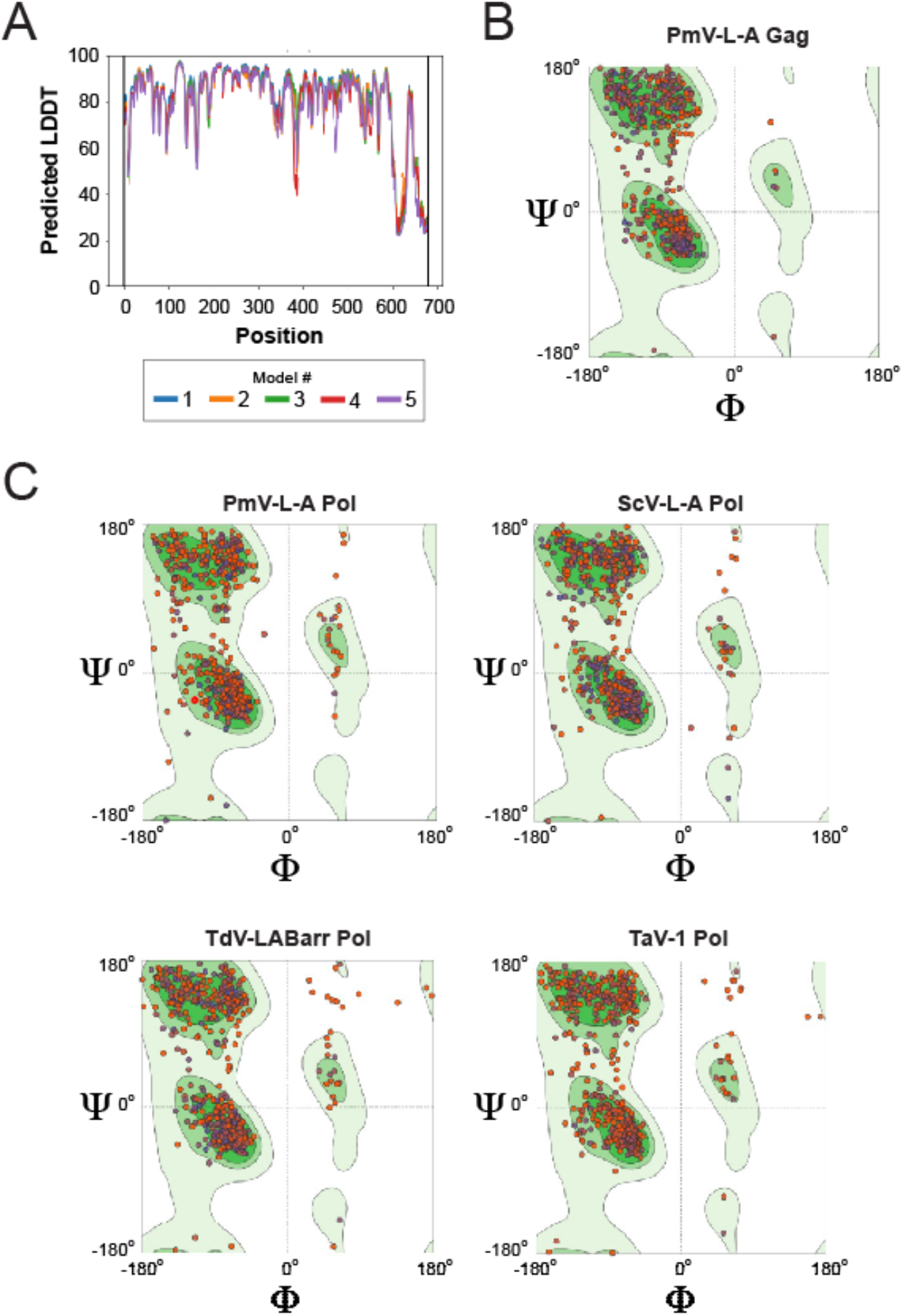
Assessment of energy minimized 3-D predicted models of Gag and Pol proteins. (A) Local distance difference test (LDDT) per residue of the AlphaFold2 predicted Gag models. Ramachandran plots post-energy minimization for each AlphaFold2 predicted model for totivirus (B) Gag and (C) four different Pol proteins. Each point represents an amino acid residue and its φ and Ψ bond angles. Shaded area of the plots represents standard angles for alpha helices, beta sheets, and left handed helices from a database of 12,521 non-redundant experimental structures.

**Table S1.** The names ans accession numbers of viruses used in this study.

| Virus | Abbreviation | Accession Number |
| --- | --- | --- |
| Saccharomyces cerevisiae virus L-BC | ScV-L-BC | NP_042580.1 |
| Scheffersomyces stipitis), | Ss_KAG2734162.1 | KAG2734162.1 |
| Tuber aestivum virus 1 | TAV-1 | YP_009507832.1 |
| Torulaspora delbrueckii virus LA | TdV-L-A | QRN76667.1 |
| Saccharomyces cerevisiae virus L-A | ScV-L-A | NP_620494.1 |
| Saccharomyces cerevisiae virus L-A-lus | ScV-L-A-lus | AFQ55380.1 |
| Saccharomyces paradoxus virus L-A-45 | SpV-L-A-45 | ATL63181.1 |
| Saccharomyces kudriavzevii virus L-A1 | SkV-L-A1 | YP_009328932.1 |
| Saccharomyces paradoxus virus L-A-62 | SpV-L-A-62 | ATL63191.1 |
| Saccharomyces paradoxus virus L-A-74 | SpV-L-A-74 | ATL63183.1 |
| Saccharomyces uvarum virus L-A-10560 | SuV-L-A-10560 | ATL63193.1 |
| Saccharomyces cerevisiae virus L-A-28 | ScV-L-A-28 | AMV49326.1 |
| Saccharomyces paradoxus virus L-A-21 | SpV-L-A-21 | ATL63179.1 |
| Red clover powdery mildew-associated totivirus 1 | RPaTV1 | BAT62477.1 |
| Maize associated totivirus | MATV | QJD25926.1 |
| Puccinia striiformis totivirus 5 | PsV5 | ATO91014.1 |
| Maize-associated totivirus 3 | MATV3 | AUH27277.1 |
| Black raspberry virus F | BRVF | YP_001497150.1 |
| Peach-associated virus 2 | PAV2 | QSV39139.1 |
| Taro-associated totivirus L | TATL | QFS21890.1 |
| Panax notoginseng virus B | PnV-A | YP_009551936.1 |
| Ambrosiozyma totivirus A | AkV-A | MK231133.1 |

**Table S2.** Confidence data for energy minimized models. Molprobity score estimates accuracy of the models in angstroms. Clashscore estimates unusual steric interaction of residues. Ramachandran favored is the percentage of residues out of the total that have acceptable angles based on standard Ramachandran angles. Ramachandran and rotamer outliers are residues that are outside of standard Ramachandran angles or have unusual rotamer conformation.

| **Model** | **Molprobrity**<br>**score** | **clashscore** | **Ramachandran**<br>**favored** | **Ramachandran**<br>**outliers** | **Rotamer**<br>**outliers** |
| --- | --- | --- | --- | --- | --- |
| PmV Gag | 1.02 | 0 | 95% | 3 | 9 |
| PmV Pol | 1.25 | 0.15 | 92.18% | 7 | 13 |
| ScV-L-A Pol | 1.37 | 0.15 | 92.06% | 5 | 19 |
| TdV-LABarr1 Pol | 1.38 | 0.82 | 90.20% | 8 | 9 |
| TAV1 Pol | 1.24 | 0.22 | 92.89 | 6 | 13 |

**File S1. PDB coordinates for the molecular models of the Gag and Pol proteins from yeast totivirues.**

## References

1. Kurtzman, C.P. Pichia E.C. Hansen (1904). In The Yeast: A Taxonomic Study; Kurtzman, C.P., Fell, J.W., Boekhout, T., Eds.; 2011; pp. 685–707 ISBN 9780080931272.

2. Zhang, J.; Xie, J.; Zhou, Y.; Deng, L.; Yao, S.; Zeng, K. Inhibitory Effect of Pichia Membranaefaciens and Kloeckera Apiculata against Monilinia Fructicola and Their Biocontrol Ability of Brown Rot in Postharvest Plum. Biol Control 2017, 114, 51–58, doi:10.1016/j.biocontrol.2017.07.013.

3. Santos, A.; Mauro, M.S.; Bravo, E.; Marquina, D. PMKT2, a New Killer Toxin from Pichia Membranifaciens, and Its Promising Biotechnological Properties for Control of the Spoilage Yeast Brettanomyces Bruxellensis. Microbiology+ 2009, 155, 624–634, doi:10.1099/mic.0.023663-0.

4. Santos, A. Killer Toxin of Pichia Membranifaciens and Its Possible Use as a Biocontrol Agent against Grey Mould Disease of Grapevine. Microbiology (Reading) 2004, 150, 2527–2534, doi:10.1099/mic.0.27071-0.

5. Melchor, R.L.A.; Rosales, V.G.; Pérez, M.C.G.; Fernández, S.P.; Álvarez, G.O.; Mastache, J.M.N. Effectiveness of Carboxylic Acids from Pichia Membranifaciens against Coffee Rust. Ciência E Agrotecnologia 2017, 42, 42–50, doi:10.1590/1413-70542018421018817.

6. Xu, X.; Chan, Z.; Xu, Y.; Tian, S. Effect of Pichia Membranaefaciens Combined with Salicylic Acid on Controlling Brown Rot in Peach Fruit and the Mechanisms Involved. J Sci Food Agr 2008, 88, 1786–1793, doi:10.1002/jsfa.3281.

7. Xu, X.B.; Tian, S.P. Reducing Oxidative Stress in Sweet Cherry Fruit by Pichia Membranaefaciens: A Possible Mode of Action against Penicillium Expansum. J Appl Microbiol 2008, 105, 1170–1177, doi:10.1111/j.1365-2672.2008.03846.x.

8. Qing, F.; Shiping, T. Postharvest Biological Control of Rhizopus Rot of Nectarine Fruits by Pichia Membranefaciens. Plant Dis 2000, 84, 1212–1216, doi:10.1094/pdis.2000.84.11.1212.

9. Belda, I.; Ruiz, J.; Alonso, A.; Marquina, D.; Santos, A. The Biology of Pichia Membranifaciens Killer Toxins. In; Toxins; 2012; Vol. 9, p. 112 ISBN 3642557589.

10. Masih, E.I.; Paul, B. Secretion of β-1,3-Glucanases by the Yeast Pichia Membranifaciens and Its Possible Role in the Biocontrol of Botrytis Cinerea Causing Grey Mold Disease of the Grapevine. Current microbiology 2002, 44, 391–395, doi:10.1007/s00284-001-0011-4.

11. Zhang, H.; Du, H.; Xu, Y. Volatile Organic Compound-Mediated Antifungal Activity of Pichia Spp. and Its Effect on the Metabolic Profiles of Fermentation Communities. Appl Environ Microb 2021, 87, doi:10.1128/aem.02992-20.

12. Young, T.W.; Yagiu, M. A Comparison of the Killer Character in Different Yeasts and Its Classification. Antonie van Leeuwenhoek 1978, 44, 59–77, doi:10.1007/bf00400077.

13. Fredericks, L.R.; Lee, M.D.; Crabtree, A.M.; Boyer, J.M.; Kizer, E.A.; Taggart, N.T.; Roslund, C.R.; Hunter, S.S.; Kennedy, C.B.; Willmore, C.G.; et al. The Species-Specific Acquisition and Diversification of a K1-like Family of Killer Toxins in Budding Yeasts of the Saccharomycotina. Plos Genet 2021, 77, e1009341, doi:10.1371/journal.pgen.1009341.

14. Ghabrial, S.A.; Castón, J.R.; Jiang, D.; Nibert, M.L.; Suzuki, N. 50-plus Years of Fungal Viruses. Virology 2015, 479–480, 356–368, doi:10.1016/j.virol.2015.02.034.

15. Xie, J.; Jiang, D. New Insights into Mycoviruses and Exploration for the Biological Control of Crop Fungal Diseases. Annual Review of Phytopathology 2014, 52, 45–68, doi: 10.1146/annurev-phyto-102313-050222.

16. Nuss, D.L. Hypovirulence: Mycoviruses at the Fungal–Plant Interface. Nature reviews Microbiology 2005, 3, 632–642, doi:10.1038/nrmicro1206.

17. Tarr, P.I.; Aline, R.F.; Smiley, B.L.; Scholler, J.; Keithly, J.; Stuart, K. LR1: A Candidate RNA Virus of Leishmania. Proc National Acad Sci 1988, 85, 9572–9575, doi:10.1073/pnas.85.24.9572.

18. Goodman, R.P.; Ghabrial, S.A.; Fichorova, R.N.; Nibert, M.L. Trichomonasvirus: A New Genus of Protozoan Viruses in the Family Totiviridae. Arch Virol 2011, 156, 171–179, doi:10.1007/s00705-010-0832-8.

19. Wang, A.L.; Yang, H.M.; Shen, K.A.; Wang, C.C. Giardiavirus Double-Stranded RNA Genome Encodes a Capsid Polypeptide and a Gag-Pol-like Fusion Protein by a Translation Frameshift. Proc National Acad Sci 1993, 90, 8595–8599, doi:10.1073/pnas.90.18.8595.

20. Herring, A.J.; Bevan, E.A. Virus-Like Particles Associated with Double-Stranded-RNA Species Found in Killer and Sensitive Strains of Yeast Saccharomyces Cerevisiae. J Gen Virol 1974, 22, 387–394, doi:10.1099/0022-1317-22-3-387.

21. Huang, S.H.; Ghabrial, S.A. Organization and Expression of the Double-Stranded RNA Genome of Helminthosporium Victoriae 190S Virus, a Totivirus Infecting a Plant Pathogenic Filamentous Fungus. Proceedings of the National Academy of Sciences of the United States of America 1996, 93, 12541–12546, doi:10.1006/viro.1998.9480.

22. Shao, Q.; Jia, X.; Gao, Y.; Liu, Z.; Zhang, H.; Tan, Q.; Zhang, X.; Zhou, H.; Li, Y.; Wu, D.; et al. Cryo-EM Reveals a Previously Unrecognized Structural Protein of a DsRNA Virus Implicated in Its Extracellular Transmission. Plos Pathog 2021, 17, e1009396, doi:10.1371/journal.ppat.1009396.

23. Tang, J.; Ochoa, W.F.; Sinkovits, R.S.; Poulos, B.T.; Ghabrial, S.A.; Lightner, D.V.; Baker, T.S.; Nibert, M.L. Infectious Myonecrosis Virus Has a Totivirus-like, 120-Subunit Capsid, but with Fiber Complexes at the Fivefold Axes. Proc National Acad Sci 2008, 105, 17526–17531, doi:10.1073/pnas.0806724105.

24. Berry, E.A.; Bevan, E.A. A New Species of Double-Stranded RNA from Yeast. Nature 1972, 239, 279–280, doi:10.1038/239279a0.

25. Fujimura, T.; Esteban, R. Cap-Snatching Mechanism in Yeast L-A Double-Stranded RNA Virus. Proc Natl Acad Sci 2011, 108, 17667–17671, doi:10.1073/pnas.1111900108.

26. Dinman, J.D.; Icho, T.; Wickner, R.B. A −1 Ribosomal Frameshift in a Double-Stranded RNA Virus of Yeast Forms a Gag-Pol Fusion Protein. Proc National Acad Sci 1991, 88, 174–178, doi:10.1073/pnas.88.1.174.

27. Esteban, R.; Wickner, R.B. A Deletion Mutant of L-A Double-Stranded RNA Replicates like M1 Double-Stranded RNA. Journal of Virology 1988, 62, 1278–1285.

28. Hall, B.G. Building Phylogenetic Trees from Molecular Data with MEGA. Mol Biol Evol 2013, 30, 1229–1235, doi:10.1093/molbev/mst012.

29. Guindon, S.; Dufayard, J.-F.; Lefort, V.; Anisimova, M.; Hordijk, W.; Gascuel, O. New Algorithms and Methods to Estimate Maximum-Likelihood Phylogenies: Assessing the Performance of PhyML 3.0. Systematic Biol 2010, 59, 307–321, doi:10.1093/sysbio/syq010.

30. Jumper, J.; Evans, R.; Pritzel, A.; Green, T.; Figurnov, M.; Ronneberger, O.; Tunyasuvunakool, K.; Bates, R.; Žídek, A.; Potapenko, A.; et al. Highly Accurate Protein Structure Prediction with AlphaFold. Nature 2021, 596, 583–589, doi:10.1038/s41586-021-03819-2.

31. Patel, J.S.; Brown, C.J.; Ytreberg, F.M.; Stenkamp, D.L. Predicting Peak Spectral Sensitivities of Vertebrate Cone Visual Pigments Using Atomistic Molecular Simulations. Plos Comput Biol 2018, 14, e1005974, doi:10.1371/journal.pcbi.1005974.

32. Abraham, M.J.; Murtola, T.; Schulz, R.; Páll, S.; Smith, J.C.; Hess, B.; Lindahl, E. GROMACS: High Performance Molecular Simulations through Multi-Level Parallelism from Laptops to Supercomputers. Softwarex 2015, 1, 19–25, doi:10.1016/j.softx.2015.06.001.

33. Blanc, A.; Ribas, J.C.; Wickner, R.B. His-154 Is Involved in the Linkage of the Saccharomyces Cerevisiae LA Double-Stranded RNA Virus Gag Protein to the Cap Structure of MRNAs and Is Essential for M1 Satellite Virus Expression. Molecular and Cellular Biology 1994.

34. Fujimura, T.; Esteban, R. Recognition of RNA Encapsidation Signal by the Yeast L-A Double-Stranded RNA Virus. J Biol Chem 2000, 275, 37118–37126, doi:10.1074/jbc.m005245200.

35. Zuker, M. Mfold Web Server for Nucleic Acid Folding and Hybridization Prediction. Nucleic Acids Res 2003, 31, 3406–3415, doi:10.1093/nar/gkg595.

36. Shen, X.-X.; Opulente, D.A.; Kominek, J.; Zhou, X.; Steenwyk, J.L.; Buh, K.V.; Haase, M.A.B.; Wisecaver, J.H.; Wang, M.; Doering, D.T.; et al. Tempo and Mode of Genome Evolution in the Budding Yeast Subphylum. Cell 2018, 175, 1533–1545.e20, doi:10.1016/j.cell.2018.10.023.

37. Naitow, H.; Tang, J.; Canady, M.; Wickner, R.B.; Johnson, J.E. L-A Virus at 3.4 Å Resolution Reveals Particle Architecture and MRNA Decapping Mechanism. Nature structural & molecular biology 2002, 9, 725–728, doi:10.1038/nsb844.

38. Lu, X.; McDonald, S.M.; Tortorici, M.A.; Tao, Y.J.; Carpio, R.V.-D.; Nibert, M.L.; Patton, J.T.; Harrison, S.C. Mechanism for Coordinated RNA Packaging and Genome Replication by Rotavirus Polymerase VP1. Structure 2008, 16, 1678–1688, doi:10.1016/j.str.2008.09.006.

39. McDonald, S.M.; Tao, Y.J.; Patton, J.T. The Ins and Outs of Four-Tunneled Reoviridae RNA-Dependent RNA Polymerases. Curr Opin Struc Biol 2009, 19, 775–782, doi:10.1016/j.sbi.2009.10.007.

40. Drinnenberg, I.A.; Fink, G.R.; Bartel, D.P. Compatibility with Killer Explains the Rise of RNAi-Deficient Fungi. Science 2011, 333, 1592–1592, doi:10.1126/science.1209575.

41. Drinnenberg, I.A.; Weinberg, D.E.; Xie, K.T.; Mower, J.P.; Wolfe, K.H.; Fink, G.R.; Bartel, D.P. RNAi in Budding Yeast. Science 2009, 326, 544–550, doi:10.1126/science.1176945.

42. Segers, G.C.; Wezel, R. van; Zhang, X.; Hong, Y.; Nuss, D.L. Hypovirus Papain-Like Protease P29 Suppresses RNA Silencing in the Natural Fungal Host and in a Heterologous Plant System. Eukaryot Cell 2006, 5, 896–904, doi:10.1128/ec.00373-05.

43. Hammond, T.M.; Andrewski, M.D.; Roossinck, M.J.; Keller, N.P. Aspergillus Mycoviruses Are Targets and Suppressors of RNA Silencing. Eukaryot Cell 2008, 7, 350–357, doi:10.1128/ec.00356-07.

44. Kubota, K.; Ng, J.C.K. Lettuce Chlorosis Virus P23 Suppresses RNA Silencing and Induces Local Necrosis with Increased Severity at Raised Temperatures. Phytopathology 2016, 106, 653–662, doi:10.1094/phyto-09-15-0219-r.

45. Frank, A.C.; Wolfe, K.H. Evolutionary Capture of Viral and Plasmid DNA by Yeast Nuclear Chromosomes. Eukaryot Cell 2009, 8, 1521–1531, doi:10.1128/ec.00110-09.

46. Khalifa, M.E.; MacDiarmid, R.M. A Novel Totivirus Naturally Occurring in Two Different Fungal Genera. Front Microbiol 2019, 10, 2318, doi:10.3389/fmicb.2019.02318.

47. Fujimura, T.; Ribas, J.C.; Makhov, A.M.; Wickner, R.B. Pol of Gag–Pol Fusion Protein Required for Encapsidation of Viral RNA of Yeast L-A Virus. Nature 1992, 359, 746–749, doi:10.1038/359746a0.

48. Routhier, E.; Bruenn, J.A. Functions of Conserved Motifs in the RNA-Dependent RNA Polymerase of a Yeast Double-Stranded RNA Virus. J Virol 1998, 72, 4427–4429, doi:10.1128/jvi.72.5.4427-4429.1998.

49. Ribas, J.C.; Fujimura, T.; Wickner, R.B. A Cryptic RNA-Binding Domain in the Pol Region of the L-A Double-Stranded RNA Virus Gag-Pol Fusion Protein. Journal of Virology 1994, 68, 6014–6020.

50. Ribas, J.C.; Wickner, R.B. RNA-Dependent RNA Polymerase Consensus Sequence of the LA Double-Stranded RNA Virus: Definition of Essential Domains. Proc National Acad Sci 1992, 89, 2185–2189, doi:10.1073/pnas.89.6.2185.

